# Plate reader microrheology

**DOI:** 10.1101/2020.07.10.197723

**Authors:** Robert F. Hawkins, Gregg A. Duncan

## Abstract

In this work, we report the development of a simplified microrheological method that can be used to rapidly study soft materials. This approach uses fluorescence polarization and a plate reader format to measure the rotational diffusion of nanoparticles within a sample of interest. We show that this measurement is sensitive to viscosity-dependent changes in polymeric soft materials and is correlated with particle tracking microrheology, a previously validated measure of microrheology. Using these fluorescence polarization-based measurements, we describe formalism that enables reasonable estimation of viscosity in polymeric materials after accounting for length-scale dependent effects of the polymer environment on the nanoparticle rotational diffusion. The use of a plate reader format allows this approach to be higher throughput, less technically challenging, and more widely accessible than standard macro- and microrheological methods, making it available to non-experts. This approach has potential applications in academic and industry settings where conventional rheological equipment may not be available, as well as in clinical settings to rapidly characterize human clinical samples.

## Introduction

Biological and synthetic polymer-based materials are integral to a wide array of industries such as in the production of adhesives, coatings, and modern medicines.^1-5^ Often, these polymeric materials exhibit both viscous and elastic properties that are important to their manufacturing and use in these applications.^6,7^ Conventional bulk rheological techniques determine the viscoelastic properties of a material by measuring its response to externally applied forces.^8,9^ However, the standard methods require a large sample volume (∼ mL), equipment that is often only accessible to highly-trained users (e.g. rheometer), and is low-throughput.^9^ Addressing some of these limitations are microrheological techniques, such as particle tracking microrheology (PTM), which can be used to determine the rheological properties of soft materials by monitoring the displacement of micrometer or nanometer-sized particles in response to thermal or applied forces.^8,9^ Microrheological methods only require a small sample volume (∼ 20 μL) which is advantageous for biological soft materials, such as peptides, DNA, and patient-derived clinical samples, that are often difficult to acquire in large volumes. However, PTM remains fairly low-throughput and uses methods that are similarly inaccessible to non-experts.^8,9^

A recent advancement in the field of microrheology is the development of techniques capable of measuring the rotational diffusion of micro-and nanoparticles, rather than their translational diffusion, to perform similar analyses as captured using PTM. For example, a technique was developed to measure microrheology based on the rotational diffusion of optically-trapped nanoparticles. As compared to standard microrheological techniques, this approach lowered the minimum sample volume needed. To address the issue of accessibility, a fully-automated dynamic light scattering (DLS) method has been used to perform rotational diffusion-based microrheological analysis on significantly lower sample volumes (down to nL) and over a greater range of frequencies compared to conventional techniques.^10,11^ However, microrheological techniques remain technically challenging to perform in a high-throughput format. This motivates our efforts to develop a simplified microrheological method that requires small sample volume, is high-throughput, and utilizes technology that is widely accessible.

Towards this end, we have developed a fluorescence polarization-based measurement that is able to produce reasonable estimates of viscosity in soft materials by measuring the rotational diffusion of fluorescent nanoparticles (NP) within a sample of interest. Motivated by the use of fluorescence polarization (FP) in many immunoassays^12-15^, we pursued this measurement approach as it can be performed in a plate reader format, making it both widely accessible and high-throughput.^16^ In this work, we investigated the sensitivity of FP-based NP rotational diffusion measurements to changes in the viscosity of various polymeric soft materials and the correlation between NP rotational diffusion, as measured by FP, and the NP translational diffusion, as measured by conventional PTM. By accounting for NP-polymer interactions, we have developed a framework to estimate viscosity in soft materials using FP for simplified plate reader microrheology.

## Theory

### Fluorescence Polarization

In FP, plane-polarized light is used to preferentially excite fluorophores that are oriented in the same direction as the polarized light.^13,17^ As the fluorophore is in its excited state, a length of time that depends on the fluorophore that is being used but is typically on the order of nanoseconds, the fluorophore may rotate due to simple Brownian motion. If the fluorophore has rotated during its excited state then the light it emits will no longer be of the same orientation as the light it was excited with – the light will be depolarized.^13,17^ The less the fluorophore rotates during its excited state, the less the light that it emits will be depolarized. Importantly, the rotational NP diffusion, as measured by the amount of depolarized light that they emit, is dependent upon the viscosity of environment, according to the Stokes-Einstein relationship (**Eq. 1**) where, *D*_r_ is the rotational diffusion coefficient, *k*_B_ is the Boltzmann constant, *T* is the temperature, *η* is the viscosity of the environment, and *R* is the radius of the spherical probe.^18^

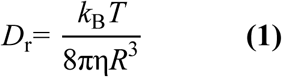

The light emitted from the fluorescent NP is passed through two filters, one parallel to the direction of the excitatory polarized light and one perpendicular, to determine the amounts of light, as measured by their intensities, that have remained polarized or become depolarized, respectively.^13,17^ Polarization (*P*) is calculated as,

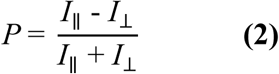

where *I*_∥_ is the intensity of emitted light that is in the same direction as the applied polarized light, *I*_⊥_ is the intensity of emitted light that is perpendicular to the direction of the applied polarized light, and *P* is the calculated polarization value.^13,17^ Therefore, a larger polarization value indicates that more light has remained polarized and the fluorescent NP has rotated less, whereas a smaller polarization value indicates that more light has been depolarized and the fluorescent NP has rotated more. Size and shape of the NP as well as the temperature and viscosity of its environment will impact the rotation and measured polarization.^13,16,17^

Closely related to the measure of FP is the measure of fluorescence anisotropy (FA). FA differs from FP only in that it takes into consideration the two directions perpendicular to the applied polarized light. For example, if a fluorophore is excited with light polarized in the Y-direction, then the depolarized light could be emitted along either the X- or Z-directions, while FP only considers the intensity of light emitted in one of these two perpendicular directions.^13,17^ Therefore, calculation of FA is similar to the calculation of FP, as can be seen in **Eq. 3**, where *r* is the calculated anisotropy value.

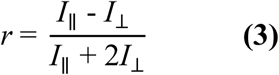

In the present work, measured *P* values are typically used in the observation of rotational diffusion. In order to normalize to the total amount of light, *P* values must be converted to *r* to perform microrheological calculations.^17^ Measured *P* values can easily be converted to *r* using **Eq. 4**.

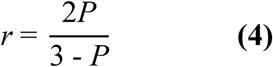

### Estimating Microviscosity

To our knowledge, the use of FP to estimate the viscosity of complex fluids has not been demonstrated. However, previous literature describes the use of FP to determine the fluidity of lipid membranes.^19,20^ In this work, the microviscosity is obtained using **Eq. 5** where *r* is the measured anisotropy, *r*_0_ is the intrinsic anisotropy that represents the value of anisotropy that would be expected if the fluorophore undergoes no rotation, *C(r)* is a parameter that takes into account molecular shape of the fluorophore as it rotates, *T* is the temperature in Kelvins, *τ* is the lifetime of the fluorophore, and *η* is the microviscosity of the environment.^20^

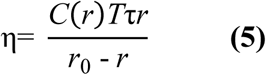

## Methods

### Nanoparticle preparation

Carboxylate modified red fluorescent polystyrene spheres with 100 nm and 500 nm diameter (Life Technologies; Excitation 530 nm & Emission 590 nm) were coated with a high surface density of polyethylene glycol (PEG) through carboxyl-amine linkage of a 5-kDa methoxy PEG-amine (Creative PEGworks) using previously established methods.^21^ Near neutral zeta potential (0 to ±5 mV) and uniform size (polydispersity index less than 0.2) of PEG-coated polystyrene NP were confirmed using a Malvern Zetasizer Nano ZS90.

### Test material preparation

Glycerol and a 20-kDa PEG (PEG20k) were mixed with UltraPure water to the desired w/w concentrations. Hyaluronic Acid (HA) and Matrigel were mixed to the desired w/v concentrations using phosphate buffered saline (PBS) with Ca^2+^ and Mg^2+^. All solutions were vortexed for 1 minute to ensure proper mixing. Matrigel was allowed to form at room temperature for 1 hour after addition of NP prior to FP and PTM measurements.

### Plate reader microrheology (PRM)

As illustrated in **Figure 1**, we have developed a new process to measure the rotational diffusion of NP using FP to determine material viscosity using PRM. Samples were prepared by pipetting the desired material (glycerol, PEG20k, Matrigel, or HA) into six wells (50 µL per well) of a black 96-well plate (Corning). Then, 0.5 µL of 10x dilute (∼0.02% w/v) 100 nm PEG-coated NP or 0.5 µL of 50x dilute (∼0.004% w/v) 500 nm PEG-coated NP were pipetted into three of the six wells of sample. Of note, NP are concentration-matched across all samples tested. After the addition of NP, the samples were allowed to equilibrate for at least 30 minutes at room temperature before FP was performed. FP was performed using a BioTek Synergy Neo2 Hybrid Multi-Mode Microplate Reader at room temperature with Excitation/Emission wavelengths of 530/590 nm, a gain of 115, and a setting of 100 reads per well. The signal intensity of the wells with NP was verified to be at least 5x greater than the signal intensity of the wells without NP. Two measurements were performed for each plate. The background signal from the samples was accounted for by subtracting the average *I*_∥_ and *I*_⊥_ of the samples without NP from the *I*_∥_ and *I*_⊥_ of each sample. Wells with negative resultant polarization values after background signal subtraction were not included for analysis. Microviscosity can be calculated using **Eq. 4** to determine *r* and **Eq. 5** to determine η.

**Figure 1.**
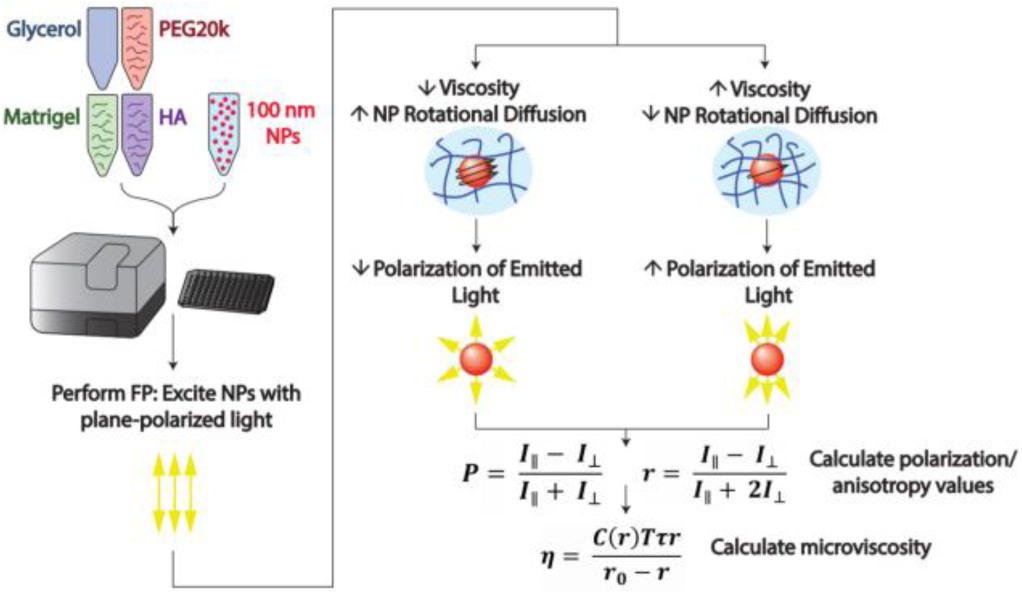
Plate reader microrheology workflow. The material test sample is combined with a dilute concentration of PEG-coated 100 nm NP into a black 96-well plate. FP is performed using a commercial multimodal plate reader. The amount of polarized and depolarized light the NP emit is used to calculate the NP rotational diffusion through the calculation of polarization and anisotropy values. The calculated anisotropy values are used to estimate the microviscosity of the environment.

### Particle tracking microrheology (PTM)

The translational diffusion of NP was measured by performing PTM using fluorescent video microscopy. Samples were prepared by adding 0.5 µL of 100x dilute (∼0.002% w/v) 100 nm PEG-coated NP or 0.5 µL of 50x dilute (∼0.004% w/v) 500 nm PEG-coated NP to 23.5 µL of sample placed within a rubber o-ring sealed to a microscope slide using vacuum grease. After the addition of NP, the samples were allowed to equilibrate for at least 30 minutes at room temperature before fluorescent video microscopy was performed. Fluorescent video microscopy was performed using a Zeiss 800 LSM microscope. Images were collected at a frame rate of 33.33 Hz for 10 seconds at room temperature. A previously developed MATLAB script was used to track the trajectories of individual NP to acquire time-averaged mean square displacement (MSD) as a function of lag time, *τ*.^22^ The generalized Stokes-Einstein relation was then used to calculate the complex modulus *G*^*\**^, as previously described.^23^ The complex viscosity, as measured by the complex modulus, is calculated as *G*^*\**^(*ω*) = *G’*(*ω*) + G*”*(*iω*) where *G’* is the storage modulus, *G’’* is the loss modulus, *i* is a complex number, and *ω* is frequency.^23^ For our purposes, the complex viscosity was calculated as the average of *G\** across frequencies 0.5-2 Hz. The pore size of the polymeric materials was determined using the equation 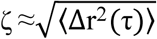, where ⟨Δr2(τ)⟩ represents the mean MSD of NP at a lag time of 1 second.^24^

## Statistical Analysis

Correlation between NP fluorescence polarization values and viscosity, and between NP fluorescence polarization values and NP translational diffusion, as measured by Pearson’s r, was calculated using GraphPad Prism. Statistical significance was considered as having a p-value < 0.05.

## Results

### Can FP measure viscosity-dependent changes in NP rotational diffusion

We hypothesized that the rotational diffusion of NP would decrease as the viscosity of the material increased. As predicted with the Stokes-Einstein equation, rotational diffusion of a spherical molecule is inversely proportional to the viscosity of the environment.^18^ Previous studies of measuring NP rotation using FP were focused on determining an unknown NP size, rather than the viscosity of the environment, and involved the development of custom optical systems.^25,26^ However, it was unknown if performing FP in a simple plate reader format would be sensitive enough to detect the viscosity-dependent changes in the NP rotational diffusion.

To test this hypothesis, we performed FP using 100 nm NP in varying concentrations of glycerol, 20-kDa polyethylene glycol (PEG20k), Matrigel, and hyaluronic acid (HA). These NP were coated in a dense layer of polyethylene glycol (PEG) to neutralize their charge in order to avoid interactions with the charged polymers of Matrigel and HA. Glycerol was chosen as a simple fluid for initial proof of concept, while the PEG20k, Matrigel, and HA were chosen due to their polymeric properties. While PEG20k and HA were prepared as polymer solutions, Matrigel forms a hydrogel containing a heterogenous mixture of collagen, laminin, and other extracellular matrix (ECM) components. The viscosity values for glycerol concentrations were acquired from literature^27^, while the viscosity values for PEG20k, Matrigel, and HA concentrations were acquired by performing PTM using 100 nm NP. We found the polarization values of 100 nm NP to be positively correlated with viscosity in all four materials tested, but only significantly so in glycerol, PEG20k, and Matrigel (**Fig. 2A-C**). The correlation between polarization values and viscosity is found to be significant in HA if only viscosities above 150 cP are considered (**Fig. 2D**). The viscosities of the tested materials ranged from approximately 1-5000 cP. We also found 100 nm NP exhibited greater sensitivity to changes in viscosity than 500 nm NP as measured by FP (**Supplemental Fig. 1**).

**Figure 2.**
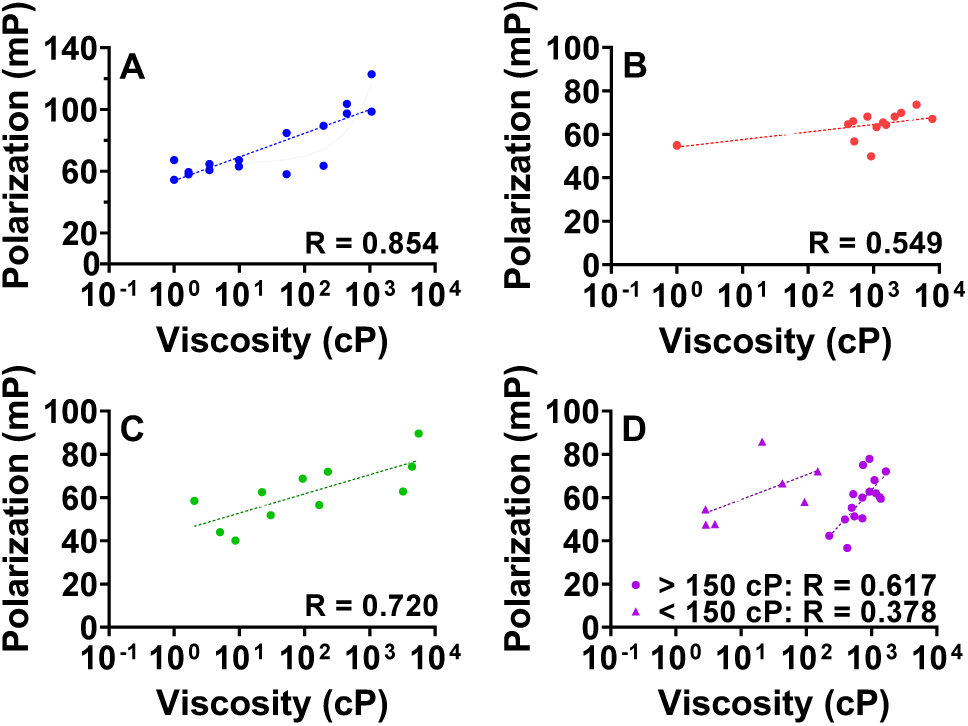
Measuring polarization of fluorescent nanoparticles in materials with varying viscosity. Measured polarization and viscosity for 100 nm NP in (**A**) glycerol [Pearson’s r = 0.854, p < 0.001], (**B**) PEG20k [Pearson’s r = 0.549, p < 0.05], (**C**) Matrigel [Pearson’s r = 0.720, p < 0.01], and (**D**) HA [<150 cP: Pearson’s r = 0.378, p > 0.05; >150 cP: Pearson’s r = 0.617, p < 0.01].

### Comparing PRM and PTM measurements

Next, we investigated the relationship between the rotational diffusion of 100 nm NP, as measured by FP, and their translational diffusion, in terms of the ensemble averaged mean-squared displacement at 1s (MSD_1s_) as measured by PTM. Since the translational diffusion of NP is a previously established and validated measure of microrheology, showing a correlation between the rotational diffusion of the NP and their translational diffusion would provide evidence that the rotational diffusion of the NP is sensitive to some of the same rheological factors and that this sensitivity is able to be detected by FP. We hypothesized that as MSD_1s_ of the NP decreased, the polarization values would increase, which would denote that the NP are rotating more slowly as they are also translating more slowly. For the 100 nm NP, we found the polarization values to be negatively correlated with the MSD_1s_ in all four materials tested and with statistical significance in glycerol, PEG20k, and Matrigel (**Fig. 3A-C**). Similarly to **Fig. 2D**, we found the correlation between FP and MSD_1s_ is only significant in HA for viscosities greater than 150 cP (**Fig. 3D**).

**Figure 3.**
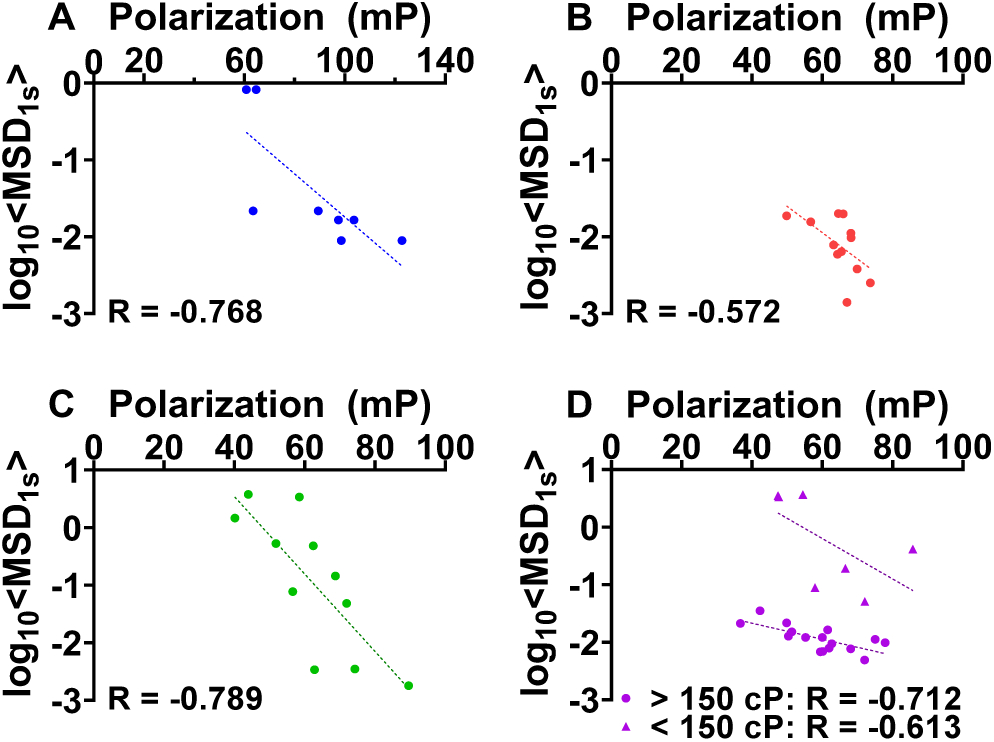
Comparing plate reader microrheology (PRM) and particle tracking microrheology (PTM) measurements. The correlation between 100 nm NP polarization values measured by PRM and log based 10 of mean squared displacement after 1 second (<log_10_MSD1s>) measured by PTM for 100 nm NP in (**A**) glycerol [Pearson’s r = -0.768, p < 0.01], (**B**) PEG20k [Pearson’s r = -0.572, p < 0.05], (**C**) Matrigel [Pearson’s r = -0.789, p < 0.01], and (**D**) HA [<150 cP: Pearson’s r = -0.613, p > 0.05; >150 cP, Pearson’s r = -0.712, p < 0.001].

### Estimating viscosity using PRM

With these initial observations to support our hypothesis, formalism was developed to interpret our measurements quantitatively and to estimate viscosity using PRM. Glycerol was chosen as the first material to estimate viscosity in due to it being a well-characterized Newtonian fluid. In order to estimate viscosity using **Eq. 5**, the values for the intrinsic anisotropy, *r*_0_, and the leading coefficients *C(r)Tτ* must be determined. Since the value for *r*_0_ is required in the determination of *C(r)Tτ, r*_0_ was determined first by measuring the polarization of 100 nm NP in 0-99.5% w/w glycerol and producing a Perrin-Weber plot in order to extrapolate to the polarization value at infinite viscosity.^13,28,29^ After fitting a linear trendline to the linear portion of the Perrin-Weber plot to determine the y-intercept, it was determined that the intrinsic polarization value, *P*_0,_ is equal to 0.124 P (P = Polarization units) for the 100 nm NP (**Supplemental Fig. 2**). This intrinsic polarization value can be converted to an intrinsic anisotropy value using **Eq. 4**, yielding a *r*_0_ value of 0.086. After *r*_0_ has been calculated, the value for the combined term of *C(r)Tτ* was determined by measuring the polarization values of 100 nm NP in 80% w/w glycerol at temperatures of 25, 27, 30, 32, and 35°C. After plotting (*r*_0_/*r* – 1)^-1^ vs. *η* and fitting a linear trendline to the produced data points, the value of *C(r)Tτ* was determined by taking the inverse of the slope of the line.^20^ This value was found to be 0.104 poise (**Supplemental Fig. 3**). When these values for *r*_0_ and *C(r)Tτ* are used in **Eq. 5**, along with the measured anisotropy values of 100 nm NP in 0-99.5% w/w glycerol, to produce PRM-based estimations of viscosity, we find generally poor agreement between these estimations and the expected viscosity values acquired from literature (**Fig. 4A**). The viscosity estimations from the 100 nm NP remained fairly constant across all of the glycerol concentrations tested, even though the expected viscosity increased by a factor of 10^3^.

**Figure 4.**
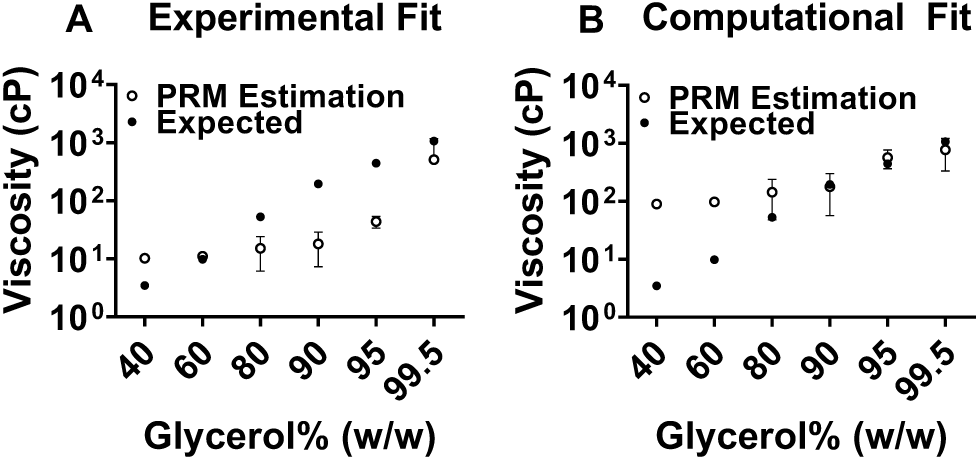
Accuracy of PRM microviscosity estimations by adjusting *C(r)Tτ* and *r*_0_ values. (**A**) PRM microviscosity estimations compared to expected microviscosity values when using experimentally determined values for *C(r)Tτ* and *r*_0_. (**B**) PRM microviscosity estimations compared to expected microviscosity values when using computationally determined values for *C(r)Tτ* and *r*_0_.

In order to address this, a procedure was created that treats the variables of *r*_0_ and *C(r)Tτ* as fitting parameters and takes as input the measured anisotropy values of the 100 nm NP in varying concentrations of glycerol to find values for *r*_0_ and *C(r)Tτ* that give the best agreement with the expected viscosity. Doing so yields values of *P*_0_ = 0.1144 P (*r*_0_ = 0.0793) and *C(r)Tτ* = 0.7636 poise for 100 nm NP. Using these new computationally fitted values for *r*_0_ and *C(r)Tτ* yields estimations of viscosity that continue to overestimate the actual viscosity in low ranges (1-100 cP), but that are more accurate within the higher range of viscosities (100-2000 cP) (**Fig. 4B**). Since using computationally fitted values for *r*_0_ and *C(r)Tτ* improved our estimations of viscosity for samples in which the expected viscosity is greater than 100 cP (**Fig. 4B**), we used this approach for our subsequent analyses.

### Estimating Viscosity of Polymer Solutions and Hydrogels using PRM

After showing the ability to produce reasonable estimations of viscosity in glycerol using PRM, we performed PRM in more complex polymer solutions of PEG20k and HA and in a polymer hydrogel of Matrigel. To benchmark PRM against the well-established PTM approach, microviscosity was determined using both approaches in PEG20k, Matrigel, and HA. For each material tested, a small volume of the sample was placed in a 96-well plate with NP to measure FP and a second small volume was placed in a microscope slide with NP to measure the NP translational diffusion via PTM. Using the fitting procedure based on measurements in glycerol, we observed underestimation of viscosity in the polymeric materials (**Supplemental Fig. 4A-D**). The cause of this underestimation was noted to be large variation in polarization values between materials with similar viscosity based on PTM measurements. For example, we see that at a given viscosity of 1000 cP the polarization values of 100 nm NP are the highest in glycerol, and then become increasingly lower in Matrigel, PEG20k, and HA (**Fig. 5A**). The lower polarization values in Matrigel, PEG20k, and HA when compared to glycerol denote that the 100 nm NP are rotating more than would be expected at the given viscosity in the polymeric materials. This increased rotation, and associated decreased polarization values, in polymeric materials leads to the underestimation of viscosity in these materials (**Supplemental Fig. 4A-D**). This greater than expected rotational diffusion appears to occur when the NP are smaller than the pore size of the material, as we see less deviation in FP values across cases when using 500 nm NP, which are larger than the pore sizes of each material (**Fig. 5B** and **Supplemental Fig. 5**).

**Figure 5.**
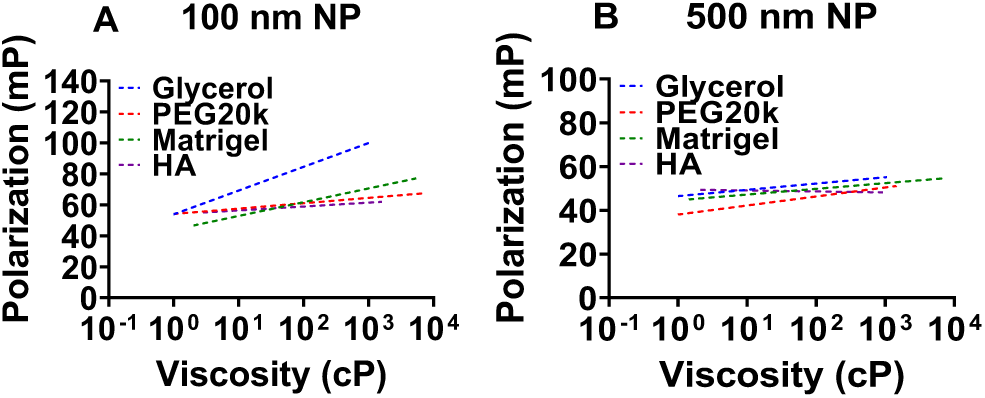
NP rotational diffusion in polymer solutions and gels. Fluorescence polarization of (**A**) 100 nm and (**B**) 500 nm NP in glycerol, PEG20k, HA and Matrigel of varying concentration and viscosity.

In past work, theory has been developed to describe length-scale dependent nanoparticle diffusion in polymeric materials.^30,31^ Maldonado-Camargo *et al*. describe that the formation of a depletion layer around the nanoparticle explains the greater than expected rotational diffusion of nanoparticles in semi-dilute polymer solutions. To account for this, a correction factor was derived based on the relationship between the nanoparticle radius, the length of the depletion layer, *δ*, and the viscosity of the polymer environment.^31^ We defined a modified form of the correction factor developed in the original work as,

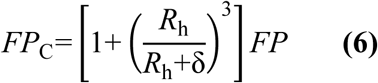

where *R*_h_ is the nanoparticle radius, *δ* is the length of the depletion layer, *FP* is the measured polarization value, and *FP*_C_ is the corrected polarization value. Meanwhile, Kohli & Mukhopadhyay describe the polymer radius of gyration, *R*_g_, as the crossover length scale for the nanoparticles to experience the viscosity of the polymer environment rather than the viscosity of the solvent. They go on to describe a correction factor based on the relationship between the nanoparticle radius, R_h_, and the polymer correlation length, ξ, that corrects for the greater than expected nanoparticle diffusion._30_ Based on this prior work, we defined a second correction factor as shown in **Eq. 7**.

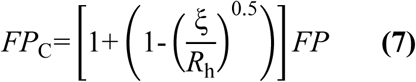

After calculating the correlation length and depletion layer length for each concentration of material tested (**Supplemental Table 1**), polarization values of the 100 nm NP follow more similar trends across all materials tested once corrected with **Eq. 6** and **Eq. 7** as shown in **Fig. 6A** and **6B**, respectively.^31-37^ After applying these correction factors, we found new values for *C(r)Tτ* and *r*_0_ of 1.67 poise and 0.096, respectively, when using the depletion layer correction factor (**Eq. 6**), and values of 1.45 poise and 0.095, respectively, when using the correlation length correction factor (**Eq. 7**) that produce more accurate estimations of viscosity across all materials (**Supplemental Fig. 4E-L**). As shown in **Fig. 6C,D**, we checked the accuracy of these correction factors by calculating the value of the correction factor that is needed to produce the desired estimation of viscosity. If the correction factor that is produced matches the correction factor that is needed, then the values should fall on to the line y = x, as denoted by the dotted line. Overall, we see reasonable clustering of the corrected measurements around the dotted line using either **Eq. 6** or **Eq. 7**.

**Figure 6.**
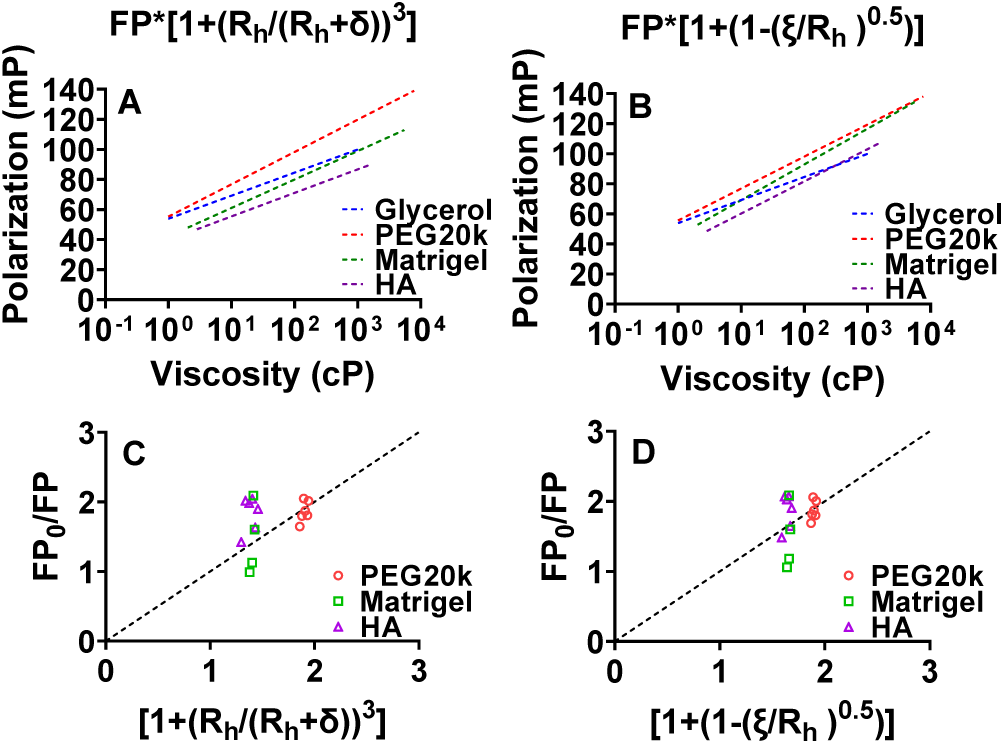
Correction factors to improve accuracy of estimated viscosity using PRM. (**A & B**) Measured polarization in glycerol, PEG20K, Matrigel, and HA versus viscosity after application of **(A)** depletion layer and **(B)** correlation length correction factors. **(C & D)** The ratio of expected (FP_0_) and measured polarization (FP) versus the calculated (**C**) depletion layer and (**D**) correlation length correction factor.

The PRM method has potential applications in academic and industry settings due to its advantages over PTM and conventional bulk rheological methods. Firstly, PRM uses a plate reader for analysis with FP as an available measurement on most commercial instruments. It is higher-throughput as multiple samples can be processed simultaneously using a plate-reader format and data analysis can be fully automated making this method approachable to non-experts. Lastly, the volume of sample needed for this approach (∼ 50 μL per well) is much less than standard rheometry where sample volumes on the order of mL are needed. The volume needed to perform PRM could be further reduced by using 96-well plates with wells of smaller size. We do not propose that this method be used to determine absolute material viscosities, but may be used as an estimation of viscosity within a reasonable range of the true value. A limitation of PRM is that it appears to have a lower limit of viscosity below which it is no longer sensitive to the sample viscosity. As noted based on measurements in HA, PRM of 100 nm NP are only significantly correlated with viscosity in HA when the viscosity of the material is above 150 cP. However, PRM remains a viable option to characterize materials used in many research and industry applications. In addition, PRM can be easily implemented into clinical laboratories to analyze the viscosity of blood, mucus, and other biological fluids.

## Conclusions

In this work, we have described a rapid and high-throughput method to estimate the viscosity of soft materials by measuring the rotational diffusion of nanoparticles using a commercially available plate reader. This method could be particularly useful to study large libraries of polymeric materials to determine optimal formulation conditions. In addition, we reason this method would be acceptable to characterize patient-derived samples of blood or mucus to determine if the sample has a normal or abnormal viscosity. Currently, this technique is limited by the range of viscosities that it is sensitive to, but this could potentially be addressed by altering the size and shape of nanoparticles used in future studies. Overall, this technique has potential to be widely used in academic, clinical, and industry settings.

## Acknowledgements

This work was supported by a Burroughs Wellcome Fund Career Award at the Scientific Interface (to GAD) and the American Lung Association.

