## Supplemental Information for "Plate reader microrheology"

### Supplemental Figures

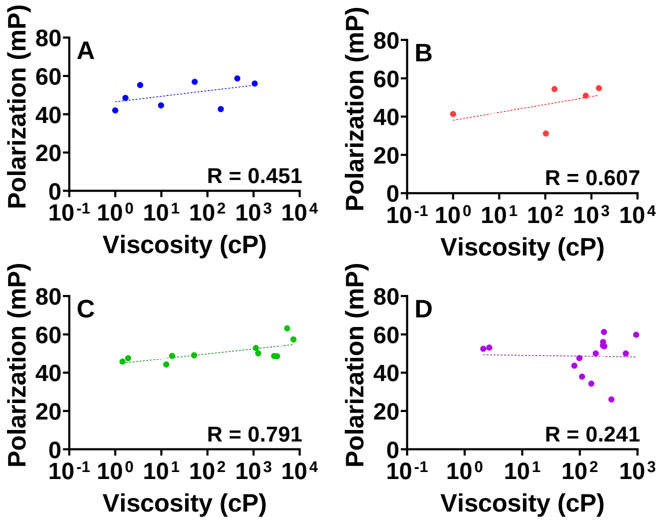


**Supplemental Figure 1. Sensitivity of rotational diffusion of 500 nm NP to changes in viscosity**. Correlation between 500 nm NP polarization values and viscosity in (**A**) glycerol [r = 0.451, p = 0.131] (**B**) PEG20k [r = 0.607, p = 0.139] (**C**) Matrigel [r = 0.791, p < 0.005] and (**D**) HA [r = 0.241, p = 0.204].

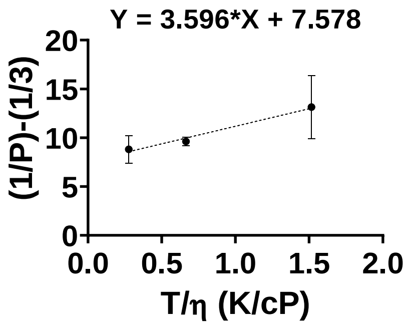


**Supplemental Figure 2. Perrin-Weber Plot to determine *$P_0$*.** After fitting a linear trendline to the produced points, the y-intercept of the line represents the expected polarization value at infinite viscosity (*$P_0$*) for 100 nm NP. Data points represent mean and standard deviation.

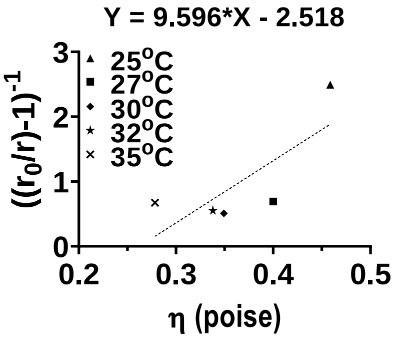


**Supplemental Figure 3. Determining the value of *$C(r)T\tau$*.** After fitting a linear trend line to the points produced by plotting of $(r_0/r - 1)^{-1}$ *vs.* $\eta$ in 80% w/w glycerol at temperatures of 25, 27, 30, 32, 35$^{o}$C, the inverse of the slope is calculated to determine the value of $C(r)T\tau$ for 100 nm NP.

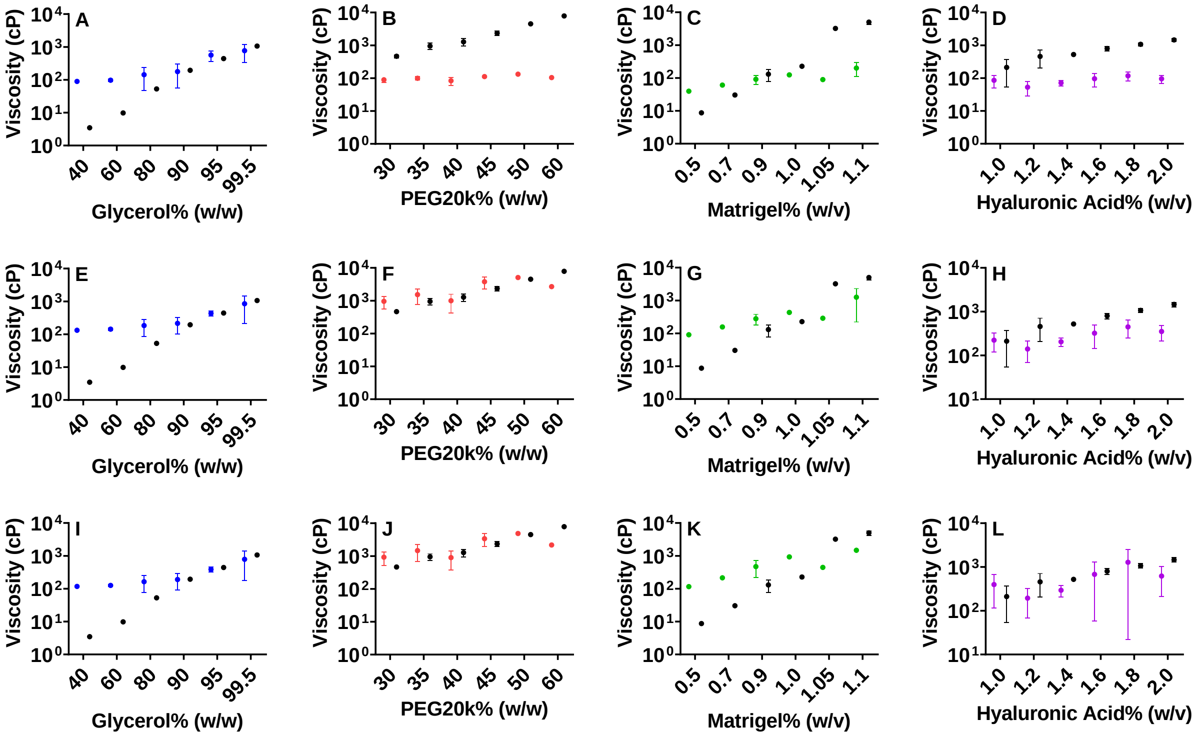


**Supplemental Figure 4. Accuracy of PRM microviscosity estimations after application of correction factors.** (**A-D**) PRM microviscosity estimations (blue, red, green, purple) compared to expected microviscosity values (black) when using computationally determined values for $C(r)T\tau$ and $r_0$ fit to glycerol in (**A**) glycerol, (**B**) PEG20k, (**C**) Matrigel, (**D**) HA. (**E-H**) PRM microviscosity estimations (blue, red, green, purple) compared to expected microviscosity values (black) after application of depletion layer correction factor and when using computationally determined values for $C(r)T\tau$ and $r_0$ fit to all materials in (**E**) glycerol, (**F**) PEG20k, (**G**) Matrigel, (**H**) HA. (**I-L**) PRM microviscosity estimations (blue, red, green, purple) compared to expected microviscosity values (black) after application of correlation length correction factor and when using computationally determined values for $C(r)T\tau$ and $r_0$ fit to all materials in (**I**) glycerol, (**J**) PEG20k, (**K**) Matrigel, (**L**) HA. Data points represent mean and standard deviation.

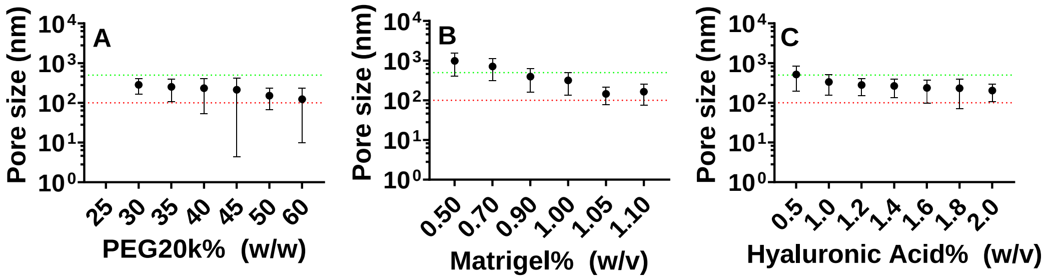


**Supplemental Figure 5. Pore size of polymeric solutions and hydrogels.** The pore size of the polymeric materials was determined using the equation $\zeta \approx \sqrt{\langle \Delta r^2(\tau) \rangle} + a$, where $\langle \Delta r^2(\tau) \rangle$ represents the mean MSD of 100 nm NP at a lag time of 1 second. Estimated pore size based on 100 nm NP MSD at 1s. (**A**) PEG20k. (**B**) Matrigel. (**C**) HA. Lines included at 100 nm (red) and 500 nm (green) as reference to size of NP used in FP-NP measurements. Data points represent mean and standard deviation.

| | ***MW* (g/mol)** | **$R_g$ (nm)** | ***c** (g/mL)** | ***c* (g/mL)** | **$\xi$ (nm)** | **$\delta$ (nm)** |
|---|---|---|---|---|---|---|
| **PEG20k** | 20000 | 6.43 | 2.98E-02 | 0.429 | 0.871 | 2.61 |
| | | | | 0.538 | 0.734 | 2.20 |
| | | | | 0.667 | 0.625 | 1.88 |
| | | | | 0.818 | 0.536 | 1.61 |
| | | | | 1.22 | 0.397 | 1.19 |
| | | | | 1.5 | 0.340 | 1.02 |
| **Matrigel** | 250000 | ~60 | 4.59E-04 | 0.005 | 10.0 | 29.4 |
| | | | | 0.007 | 7.77 | 23.0 |
| | | | | 0.009 | 6.44 | 19.2 |
| | | | | 0.01 | 5.95 | 17.7 |
| | | | | 0.0105 | 5.73 | 17.1 |
| | | | | 0.011 | 5.54 | 16.5 |
| **HA** | 500000 | 68.54 | 6.16E-04 | 0.01 | 8.47 | 25.1 |
| | | | | 0.012 | 7.39 | 21.9 |
| | | | | 0.014 | 6.58 | 19.6 |
| | | | | 0.016 | 5.95 | 17.7 |
| | | | | 0.018 | 5.45 | 16.3 |
| | | | | 0.02 | 5.04 | 15.0 |

**Supplemental Table 1. Correlation length and depletion layer length of polymeric materials.** The correlation length of each concentration used is calculated using the equation $\xi = R_g \left(\frac{c}{c^*}\right)^{-0.75}$, where $R_g$ is the radius of gyration of the material, *c* is the concentration of the material (g/mL), and $c^*$ is the critical overlap concentration of the material (g/mL). $c^*$ is calculated by the equation $c^* = \left(\frac{3MW}{4\pi R_g{}^3 N_A}\right)$, where *MW* is the molecular weight of the polymer (g/mol) and $N_A$ is Avogadro's number. The depletion layer length for each concentration used is calculated using the equation $\delta = 3\left(\frac{R_h{}^2 \xi^2}{R_h{}^2 + \xi^2}\right)$, where $R_h$ is the radius of the NP.
